# Transition of a Yeast Endosymbiont from a Free-living to Host-reliant Lifestyle Through Gene Loss and Horizontal Gene Transfer

**DOI:** 10.64898/2026.06.29.735303

**Authors:** Trina Roychoudhury, Janardhan Pallavi, Anupam Roy, Anindita Seal

## Abstract

Endosymbiosis is widespread throughout the tree of life. Understanding how the transition of a bacterial endosymbiont from facultative to host-dependent obligate life occurs is an important question for defining the origin of endosymbiosis. A novel gram-positive bacillus, *Brevibacillus* sp. TJ4 was isolated from the nitrogen-fixing yeast *Rhodotorula mucilaginosa* JGTA-S1, which houses several endobacteria within its cells. TJ4 can survive independently of yeast but exhibits genomic and metabolic features characteristic of an evolving endosymbiont, slowly assuming a host-dependent, obligate lifestyle. The TJ4 genome contains several incomplete pathways for carbohydrate, amino acid, vitamin, and cofactor metabolism, which is reflected in its increased reliance on host-derived nutrients and auxotrophy compared with that of other *Brevibacillus* spp. Comparative genomics revealed widespread genome rearrangements, loss of synteny, and multiple cross-genus and inter-kingdom horizontal gene transfer (HGT) events in TJ4 compared to other *Brevibacillus* spp. These HGTs include the acquisition of genes from bacteriophages and co-resident endobacteria of JGTA-S1. One such horizontally acquired gene, Type II 3-dehydroquinate dehydratase (AroQ), appears to have originated from the *Rhodotorula* host itself. This acquisition functionally restores the shikimate pathway in strain TJ4, as evidenced by the phylogenetic placement of AroQ from TJ4 within the clade of fungal AroQ homologs. Potential exploitation of the host JGTA-S1 appears to be a probable mode of endosymbiosis of TJ4, an evolving endosymbiont that we named *Brevibacillus rhodotorulae* sp. nov.

## Introduction

The Intimate association between a host and its endosymbiont significantly influences the niche adaptation and evolutionary trajectory of both organisms. However, the specific mechanisms and underlying reasons driving endosymbiotic colonization remain largely elusive in most systems. Although endosymbiosis is frequently perceived as a relationship that benefits the host organism, it is not always the case. In many instances, this relationship may initially be exploitative, with the endosymbiont deriving an advantage at the expense of the host. Over evolutionary timescales, such associations can shift, eventually becoming mutualistic, thereby justifying the continued presence of the endosymbiont despite the potential burden it may impose on its host.

The basidiomycetous endophytic yeast *Rhodotorula mucilaginosa* JGTA-S1 is a complex holobiont that hosts a diverse community of endosymbiotic bacteria, some of which in many ways regulate its fitness and interaction with plants, such as rice (Nag et al. 2024; Paul et al. 2020). Our previous study showed that JGTA-S1 colonizes rice plants and promotes their growth, development, and nitrogen nutrition. JGTA-S1 yeast has a unique ability to transform into a filamentous form associated with rice roots, allowing bacteria to invade these filaments (Paul et al., 2020). Shotgun whole genome metagenomic data confirmed the presence of a microbial community, leading us to adopt a culturomics strategy (Nag et al. 2024) to isolate the culturable JGTA-S1 endosymbiont, *Brevibacillus* sp. TJ4.

The physical proximity of endobacteria inside JGTA-S1 creates an ideal setting for genomic flux via horizontal gene transfer (HGT). Microbial evolution is driven by HGT, a common natural process that allows genetic material to be exchanged across species boundaries to support adaptation to specialized habitats. The environmental context often acts as a selective force, promoting HGT and conferring specific advantages to recipient bacteria. For example, in the cheese microbiome, certain bacteria acquire genes that enable enhanced iron uptake, thereby gaining a competitive edge over other microbial inhabitants (Bonham et al., 2017). The role of HGT has been thoroughly documented in complex environments, such as the mammalian gut and soil microbiomes. In contrast, fungal holobionts and associated HGT processes remain less explored, primarily because of the challenges in cultivating the interacting microbial species.

The microbiome of *R. mucilaginosa* JGTA-S1 is remarkable because of the unicellular nature of the yeast, which imposes spatial constraints and compels resident microbes to engage in close and potentially complex interactions. Different JGTA-S1 endosymbionts can be isolated and cultured, albeit with varying degrees of difficulty. Endosymbionts are often classified based on their genome reduction, transmission pattern (horizontal vs. vertical), and dependence on the host (obligate vs. facultative) (Araldi-Brondolo et al. 2017). The abundance of these endosymbiotic bacteria changes in response to nitrogen availability in yeast (Nag et al., 2024). Within the culturable community of JGTA-S1 endobacteria, *Brevibacillus* sp. TJ4 stands out as a novel bacterium whose genome exhibits hallmark traits of an ongoing shift from a free-living existence to host dependence, marking a significant proto-obligate stage in the evolution of endosymbionts. We named this new species *Brevibacillus rhodotorulae*. Therefore, JGTA-S1 and its endosymbionts represent an excellent model for studying ongoing endosymbiotic relationships.

## Results

### *Brevibacillus* sp. TJ4 is isolated from *Rhodotorula mucilaginosa* JGTA-S1

To isolate the cultivable endosymbionts of *Rhodotorula mucilaginosa* JGTA-S1, the yeast was grown in a nitrogen-free medium, ruptured, and the extract was plated on tryptic soy broth (TSB) agar plates (Nag et al. 2024). A rich medium (TSB) was used so that the majority of the endobacteria could be cultured, including those that lost genes for nutrition and metabolism during the course of becoming endosymbionts. A pure culture of a yellow-colored bacillus (Fig 1A) was obtained, and whole-genome sequencing was performed using Illumina and Oxford Nanopore platforms. The genome was analyzed using the Type Strain Genome Server (TYGS) to identify the bacterium. The genome contained two contigs, 4389 genes, 30 rRNAs, and 103 tRNAs.

**Fig 1.**
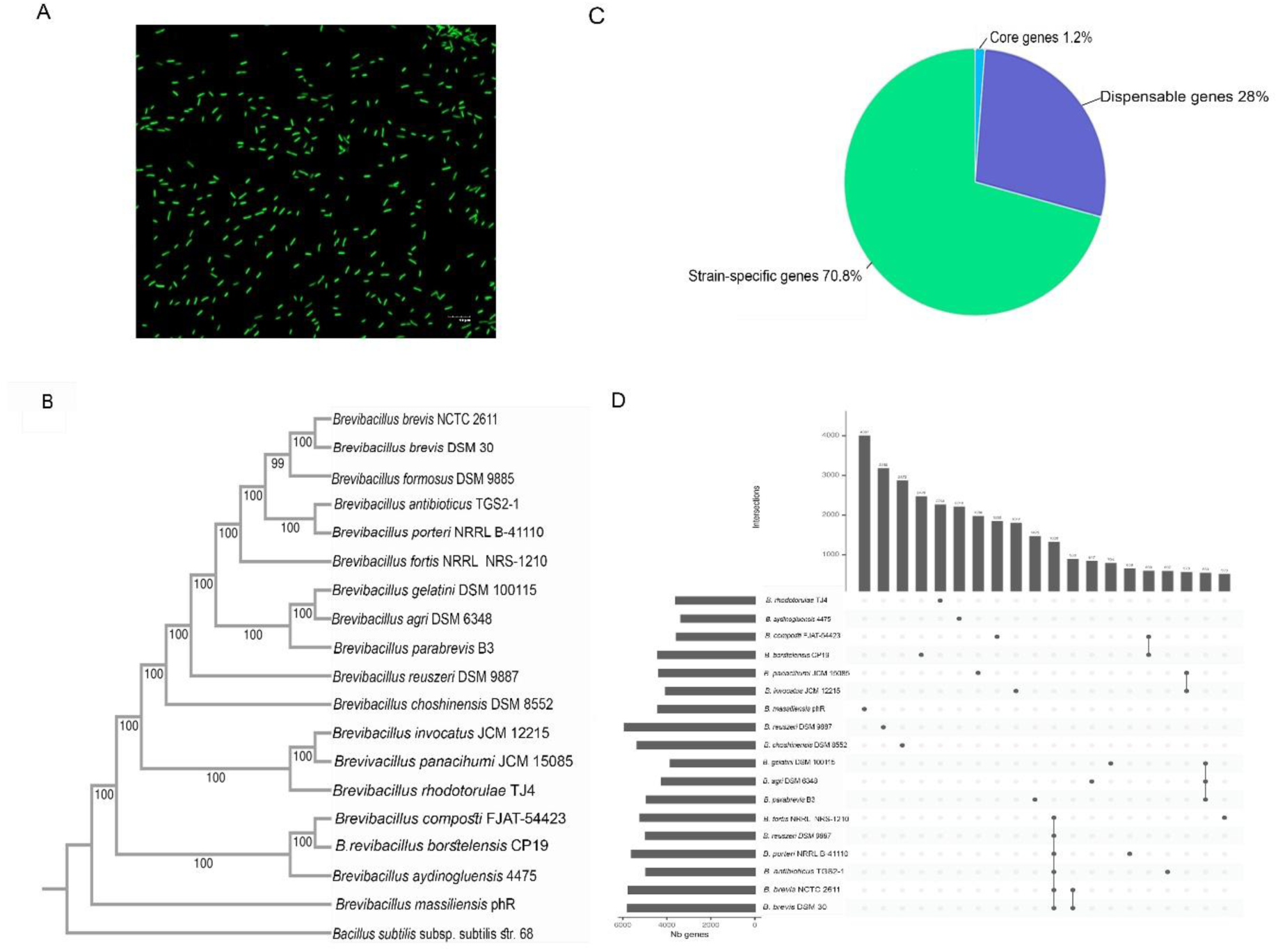
*Brevibacillus* sp. TJ4 is positioned distinctly in a phylogenetic tree with high number of species-specific unique genes. Whole genome-based phylogeny using 17 *Brevibacillus* genomes along with *Brevibacillus* sp. The TJ4 was performed using the BVBRC platform. A pie chart was generated using BLAST analysis of annotated TJ4 genes, Core and genome pan-genome analysis, are presented by an Upset plot A. Microscopic image of TJ4. B. Whole genome-based phylogenetic tree for 18 *Brevibacillus* genomes, C. Distribution of core, dispensable and strain specific genes. D. Upset Plot showing clusters of shared genes between 18 *Brevibacillus* genomes.

### *Brevibacillus* sp. TJ4 is a novel species with several lost and unique genes

To study the TJ4 genome and compare it with other *Brevibacillus* species, the hybrid assembly of TJ4 and genomes of 17 other *Brevibacillus* species were subjected to genome-based phylogenetic analysis using the BV BRC web server. The TJ4 genome was distantly segregated from its nearest neighbors in a phylogenetic tree (Fig 1B) and had only 22.9 % similarity to its closest known species according to digital DNA-DNA hybridization data from the Genome-to-Genome Distance Calculator (Table 1). This distinct clustering from its closest neighbors indicates a unique environment from which it was isolated. Among the *Brevibacillus* genomes examined, TJ4 possessed the second smallest genome after *B. aydinogluensis* 4475 (Table 1). A search was conducted to identify unique genes present in TJ4. Genome analysis revealed that core genes universally present in all tested *Brevibacillus* species constituted only 1.2%, indicating significant genetic differences among the *Brevibacillus* strains. The observed lower percentage of core and dispensable genes (genes shared between multiple *Brevibacillus* species) coupled with a high proportion of strain-specific genes suggests significant adaptability and ongoing evolutionary divergence within the *Brevibacillus* strains, driven by continuous gene gain and loss (Fig 1C) (Tettelin et al. 2005). This diversity is reflected in the UpSet plot, an alternative representation of the Venn diagram, where the intersection sizes of genes of different *Brevibacillus* strains, including TJ4, are shown (Fig 1D). For TJ4, the number of unique clusters was 2269. The TYGS server suggested that TJ4 is a novel species; hence, we propose the name *Brevibacillus rhodotorulae* sp. nov. MTCC 13946^T^ for TJ4, which is consistent with its source of isolation.

**Table 1.** *Brevibacillus* sp. TJ4 is a novel *Brevibacillus* according to DNA-DNA hybridization data.

| Query strain<br>(Genome<br>size) | Subject strain<br>(Genome size) | dDDH<br>(d0,<br>in %) | C.I.<br>(d0,<br>in %) | dDDH<br>(d4,<br>in %) | C.I.<br>(d4,<br>in %) | dDDH<br>(d6,<br>in %) | C.I.<br>(d6,<br>in %) | G+C<br>content<br>difference<br>(in %) |
| --- | --- | --- | --- | --- | --- | --- | --- | --- |
| *B. rhodotorulae*<br>TJ4 (4,348,424<br>bp) | *Brevibacillus borstelensis*<br>CP19 (5,570,859 bp) | 15.2 | [12.3 -<br>18.6%] | 22.9 | [20.6 -<br>25.4%] | 15.4 | [12.9 -<br>18.3%] | 2.11 |
|  | *Brevibacillus composti* FJAT-54423 (4,457,958 bp) | 15.1 | [12.2 -<br>18.5] | 20.7 | [18.5 -<br>23.1] | 15.2 | [12.7 -<br>18.1] | 0.59 |
|  | *Brevibacillus agri* DSM 6348<br>(5,374,818 bp) | 17.8 | [14.8 -<br>21.4%] | 20.4 | [18.2 -<br>22.8%] | 17.5 | [14.9 -<br>20.5%] | 0.13 |
|  | *Brevibacillus aydinogluensis*<br>4475 (4,105,500 bp) | 15.7 | [12.8 -<br>19.2%] | 20.3 | [18.1 -<br>22.7%] | 15.7 | [13.2 -<br>18.7%] | 2.52 |
|  | *Brevibacillus panacihumi*<br>JCM 15085 (5,116,160 bp) | 18.6 | [15.4 -<br>22.1] | 20.1 | [17.9 -<br>22.5] | 18.1 | [15.4 -<br>21.1] | 2.74 |
|  | *Brevibacillus parabrevis* B3<br>(6,088,079 bp) | 16.6 | [13.6 -<br>20.2%] | 20.1 | [17.9 -<br>22.5%] | 16.5 | [13.9 -<br>19.4%] | 1.53 |
|  | *Brevibacillus gelatini* DSM<br>100115 (4,637,568 bp) | 17.6 | [14.5 -<br>21.1] | 19.9 | [17.6 -<br>22.3] | 17.3 | [14.7 -<br>20.2] | 1.73 |
|  | *Brevibacillus antibioticus*<br>TGS2-1 (6,181,736 bp) | 15.3 | [12.4 -<br>18.7] | 19.8 | [17.6 -<br>22.2] | 15.3 | [12.8 -<br>18.2] | 6.69 |
|  | *Brevibacillus brevis*<br>NCTC2611 (6,726,998 bp) | 15.5 | [12.5 -<br>18.9%] | 19.6 | [17.4 -<br>22%] | 15.5 | [13 -<br>18.4%] | 6.29 |
|  | *Brevibacillus choshinensis*<br>DSM 8552 (6,268,382 bp) | 15.9 | [13.0 -<br>19.4] | 19.4 | [17.2 -<br>21.8] | 15.9 | [13.3 -<br>18.8] | 5.36 |
|  | *Brevibacillus invocatus* JCM<br>12215 (4,753,702 bp) | 18.6 | [15.5 -<br>22.2] | 19.2 | [17.0 -<br>21.6] | 18 | [15.4 -<br>21.0] | 4.8 |
|  | *Brevibacillus fortis* NRRL<br>NRS-1210 (6,268,222 bp) | 15.2 | [12.3 -<br>18.6] | 18.9 | [16.7 -<br>21.3] | 15.2 | [12.7 -<br>18.1] | 6.58 |
|  | *Brevibacillus reuszeri* DSM<br>9887 (6,968,040 bp) | 15.1 | [12.2 -<br>18.5] | 18.8 | [16.6 -<br>21.2] | 15.1 | [12.6 -<br>18.0] | 6.78 |
|  | *Brevibacillus formosus* DSM<br>9885 (6,212,749 bp) | 15.4 | [12.4 -<br>18.8] | 18.8 | [16.6 -<br>21.2] | 15.4 | [12.9 -<br>18.2] | 6.31 |
|  | *Brevibacillus brevis* DSM 30<br>(6,609,649 bp) | 15.3 | [12.4 -<br>18.8] | 18.7 | [16.5 -<br>21.0] | 15.3 | [12.8 -<br>18.2] | 6.35 |
|  | *Brevibacillus porteri* NRRL B-41110T (6,462,020 bp) | 15.3 | [12.4 -<br>18.7] | 18.6 | [16.4 -<br>20.9] | 15.3 | [12.8 -<br>18.2] | 6.62 |
|  | *Brevibacillus massiliensis*<br>pHR (4,952,787 bp) | 13.3 | [10.6 -<br>16.6] | 17.6 | [15.5 -<br>20.0] | 13.6 | [11.2 -<br>16.4] | 0.61 |

### *Brevibacillus rhodotorulae* has incomplete metabolic pathways for carbohydrates, amino acids, vitamins, and co-factors

The KEGG Mapper reconstruction tool was used to study the integrity of different metabolic pathways in *B. rhodotorulae*. The data revealed that the bacterium possesses many incomplete pathways, including those of modules with carbohydrates, amino acids, vitamins, and cofactor metabolism (Table S1 and Fig. 2). Analysis of the genome revealed incomplete pathway modules, indicating that several essential KEGG Orthologs (KOs) required for the assembly of fully functional metabolic and signaling systems are absent. Data were illustrated using iPath3.0 (Darzi et al. 2018) with absent genes in carbohydrate, amino acid, vitamin, and cofactor metabolism marked in IBM purple, orange, and electric blue, respectively (Fig S1). The corresponding present and missing pathways are shown in purple, orange, and cobalt blue (Fig 2). Additionally, *B*. *rhodotorulae* lacks many genes involved in methane, nitrogen, sulfur, fatty acid, lipid, pyrimidine, and other polysaccharide metabolism, nucleotide sugar, and glycan biosynthesis (Table S1).

**Fig 2.**
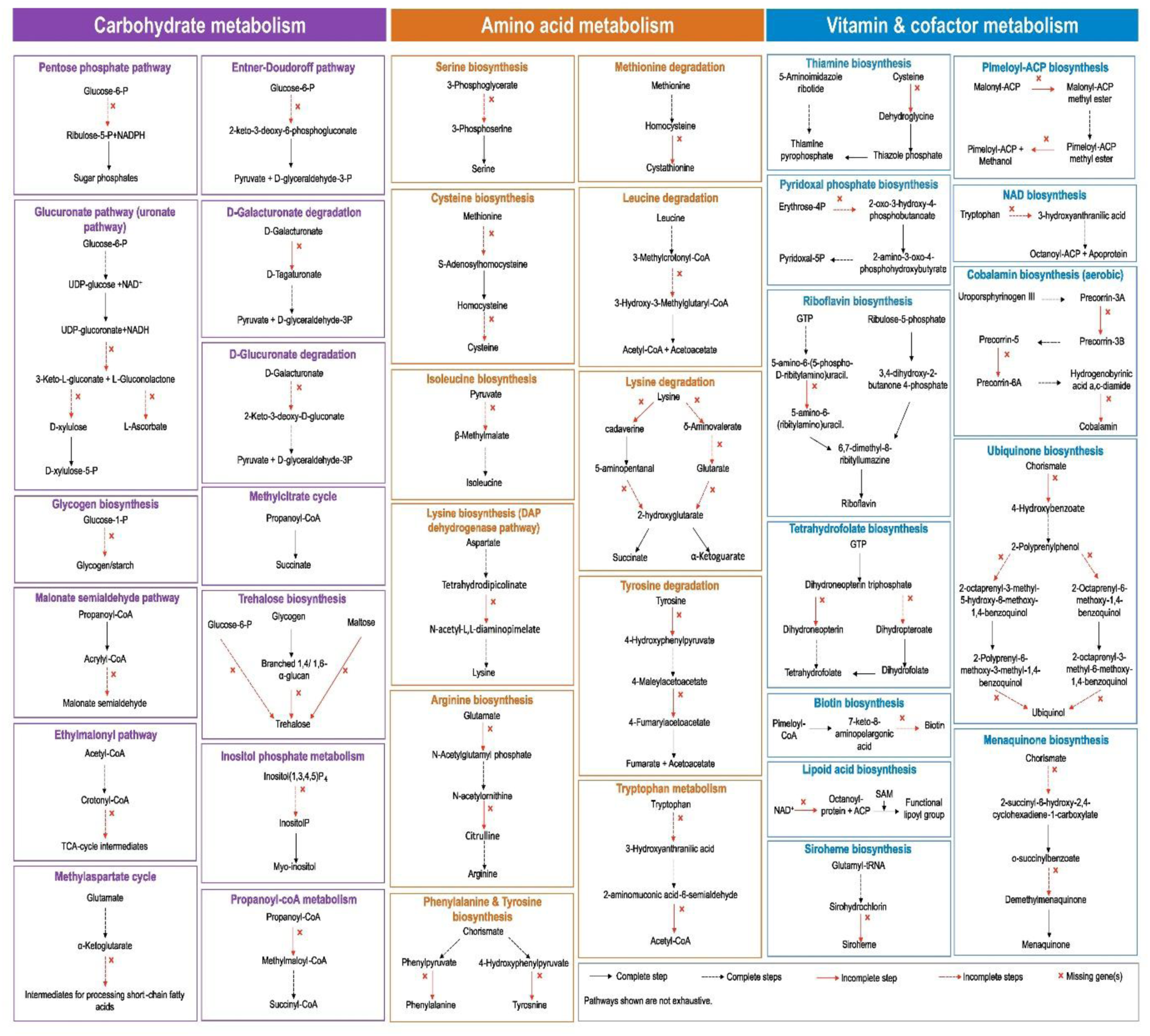
*Brevibacillus rhodotorulae* has defects in many metabolic pathways affecting its energy metabolism. KofamKOALA was used to search for KEGG orthologs of bacterial metabolic pathways in *Brevibacillus rhodotorulae*. The genes missing blocks in carbohydrate, amino acids, vitamin, and cofactor metabolic pathways are marked in purple, orange, and cobalt blue respectively. Each black solid arrow represents a complete step 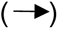, Each black dashed arrow represents multiple complete reaction steps 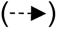, Each red solid arrow represents an incomplete step 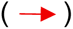, each red dashed arrow represents multiple incomplete reaction steps 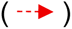, missing gene(s) 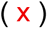.

#### Carbohydrate metabolism

*B*. *rhodotorulae* has incomplete pentose phosphate (PPP) and Entner–Doudoroff (ED) pathways. The PPP normally runs parallel to glycolysis and produces NADPH and pentose phosphates, whereas the ED pathway is an alternative route for converting glucose to pyruvate. *B*. *rhodotorulae* lacks glucose-6-phosphate dehydrogenase (G6PD), hexose-6-phosphate dehydrogenase (H6PD), 6-phosphogluconolactonase (PGLS), and 6-phosphogluconate dehydratase (EDD), indicating that both the oxidative PPP and ED pathways are inactive (Fig 2, 3A). This increases sensitivity to oxidative stress, consistent with the inability of strain TJ4 to grow in paraquat dichloride, unlike *B*. *velezensis* TSB6.1 ̶ another JGTA-S1 endosymbiont (NZ_JAOXLR000000000.1, Nag and Seal 2024), which retains an intact oxidative PPP and nearly complete ED pathway (Fig 3A). Because the ED pathway also supports the metabolism of sugar acids, such as D-galacturonate, gluconate, D-galactonate, and D-glucuronate, its loss would normally reduce metabolic flexibility under nutrient-poor conditions. However, as an endofungal bacterium, *B*. *rhodotorulae* likely compensates for this by acquiring nutrients from its fungal host. It is also deficient in trehalose biosynthesis, which is important for energy storage and for stress protection. In addition, the ethylmalonyl pathway, which supports growth on C1 and C2 compounds, is incomplete, as are the methylaspartate and methylcitrate cycles, potentially limiting carbon assimilation and causing toxic accumulation of propionyl-CoA. The malonate semialdehyde pathway (propanoyl-CoA => acetyl-CoA) is absent, likely impairing valine degradation, and the pathway converting propanoyl-CoA to succinyl-CoA is also incomplete. Finally, inositol phosphate metabolism [Ins(1,3,4,5)P4 => Ins(1,3,4)P3 => myo-inositol], which is important for carbon recovery, energy use, and lipid synthesis, is also incomplete in *B. rhodotorulae*.

**Fig 3.**
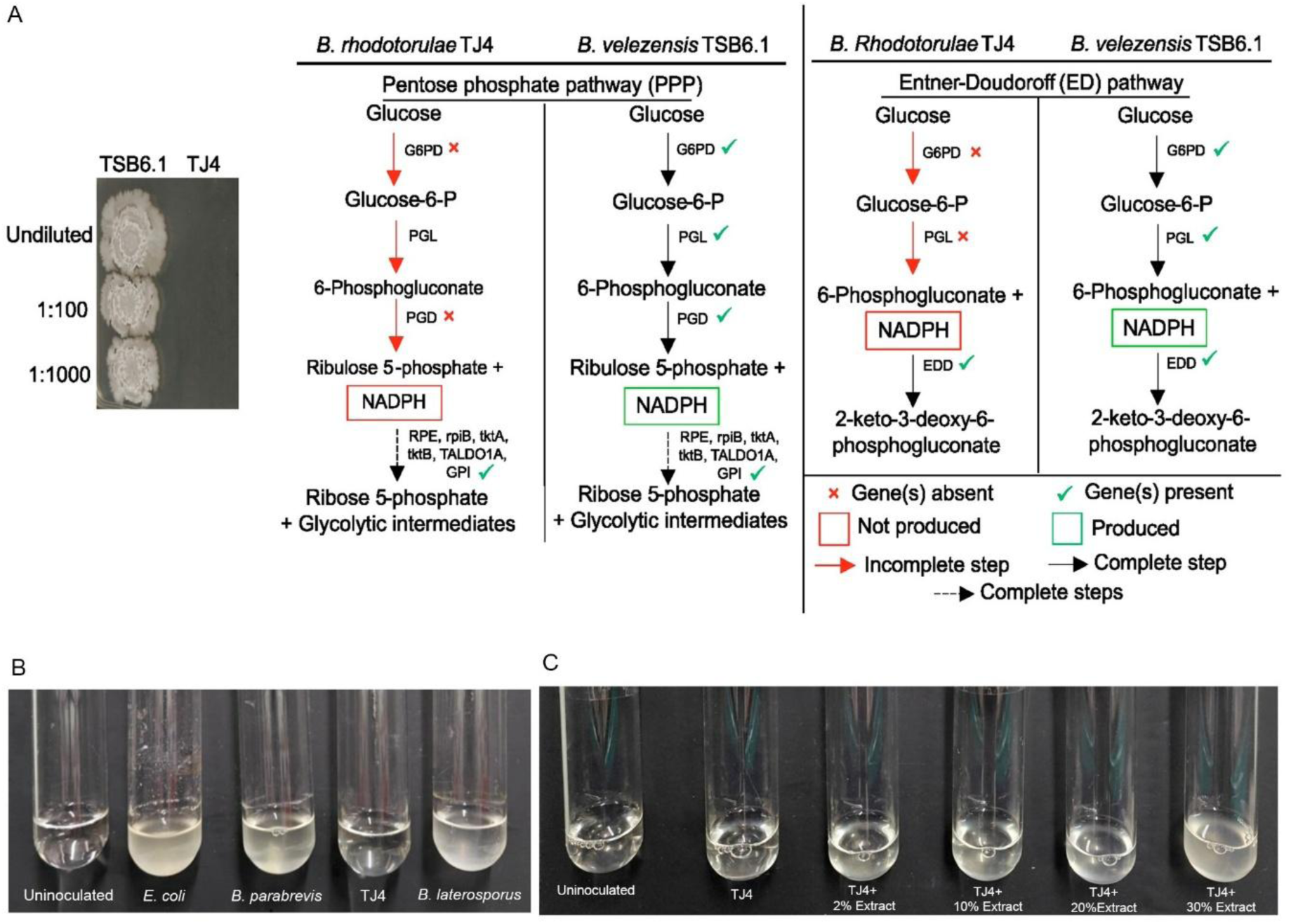
*Brevibacillus rhodotorulae* is sensitive under unfavourable conditions. *Brevibacillus rhodotorulae* was grown in TSB in the presence of 1 µM paraquat dichloride or in minimal M9 medium with or without supplementation of JGTA-S1 extracts at different concentrations A. Spotting assay of *Brevibacillus rhodotorulae* and *Bacillus velezensis* TSB6.1. Undiluted (OD_600_ 0.5), 1: 100 (OD_600_ 0.005), 1: 1000 (OD_600_ 0.0005). Comparative PPP and ED pathways of TJ4 and TSB6.1 are shown. Each black solid arrow represents a complete step 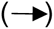, Each black dashed arrow represents multiple complete reaction steps 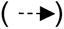, Each red solid arrow represents an incomplete step 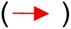, missing gene(s) 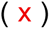, gene(s) present 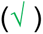, produced 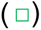, not produced 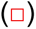, B. TJ4 was grown in M9 medium for 14 d and imaged, C. TJ4 grown in M9 medium for 3 d was supplemented with JGTA-S1 extract.

#### Amino acid metabolism

*B*. *rhodotorulae* lacks several genes required for the synthesis of different amino acids, such as cysteine (methionine => cysteine), serine (polar), arginine, lysine (positively charged), isoleucine (nonpolar), tyrosine, and phenylalanine (aromatic) (Table S1 and Fig 2). The inability to synthesize essential molecules, such as amino acids, would have serious repercussions for any organism. Additionally, this bacterium lacks different genes required for the degradation of methionine, leucine, lysine, hydroxyproline, tyrosine, and tryptophan, which would result in wider systemic problems for bacteria, such as protein synthesis interference, metabolic imbalance, energy deprivation, and accumulation of toxic intermediates.

#### Vitamin and cofactor metabolism

This bacterium cannot synthesize many essential vitamins that act as coenzymes/co-factors in many vital reactions for energy metabolism, such as pyridoxal-phosphate, pimeloyl-ACP, cobalamin (aerobic, uroporphyrinogen III => precorrin 2 => cobyrinate a,c-diamide), biotin, NAD (tryptophan => quinolinate => NAD), and tetrahydrofolate. It also cannot produce the prosthetic group siroheme, antioxidants ubiquinone, lipoic acid, and menaquinone, which are important for energy generation. This metabolic dependency on the host strongly suggests that this bacterium has an endosymbiotic lifestyle.

Such deletions may affect energy metabolism and nucleic acid and amino acid synthesis. These notions are supported by the failure of the organism to grow in minimal medium, such as M9, unlike *B. parabrevis* and *B. laterosporus*, which grew in the same medium (Fig 3B). *Escherichia coli* was used as a positive control. The growth of *B*. *rhodotorulae* was best restored by supplementation of 30% JGTA-S1 extract into the medium (Fig 3C), suggesting that the bacterium obtained critical nutrients from its fungal host. TJ4 showed slow growth even in enriched media, such as TSB (data not shown).

Additionally, *B*. *rhodotorulae* lacks many genes for sporulation (GerW family sporulation protein, YyaC-like protein, sporulation protein Spo0E, Spo0M, Stage II sporulation protein P, R, Stage III sporulation protein AF, AH, Stage V sporulation protein B, E, and YhcN/Ylaj family sporulation lipoprotein), making it susceptible to stress adaptation (Table S2). This bacterium is also deficient in GABA, a signaling molecule crucial for bacterial behavior, including spore germination, and polyamines, which are important for cell growth and stress resistance.

#### Horizontal gene transfer in *B*. *rhodotorulae*

As a novel species, *B*. *rhodotorulae* was compared with 17 other *Brevibacillus* species. This analysis identified 84 unique genes with known functions in *B*. *rhodotorulae*, excluding hypothetical, putative, and proteins with domains of unknown functions (DUFs) (Table S2, Fig 4A). The distinct phylogenetic segregation of TJ4 suggests horizontal gene transfer (HGT), a common driver of prokaryotic genome evolution. Such exchanges may be favored by the close endosymbiont-host association within JGTA-S1. To detect genomic islands and HGT events, we analyzed the genome using IslandViewer 4 and Alien Hunter (Fig 4B, Table S3). IslandViewer 4 identifies regions with atypical sequence compositions, including codon bias and GC content differences, whereas Alien Hunter detects compositional bias based on variable-order motif distributions. Only two predicted HGT genes were shared by both methods: L-arabinose isomerase (AraA) and ribulokinase (AraB_1). These genes are involved in L-arabinose degradation, which supports energy production via the pentose phosphate pathway. However, they are not unique to *B*. *rhodotorulae* and also occur among the dispensable genes in *B*. *massiliensis* phR (AraA) and *B*. *aydinogluensis* 4475 (AraB_1).

**Fig 4.**
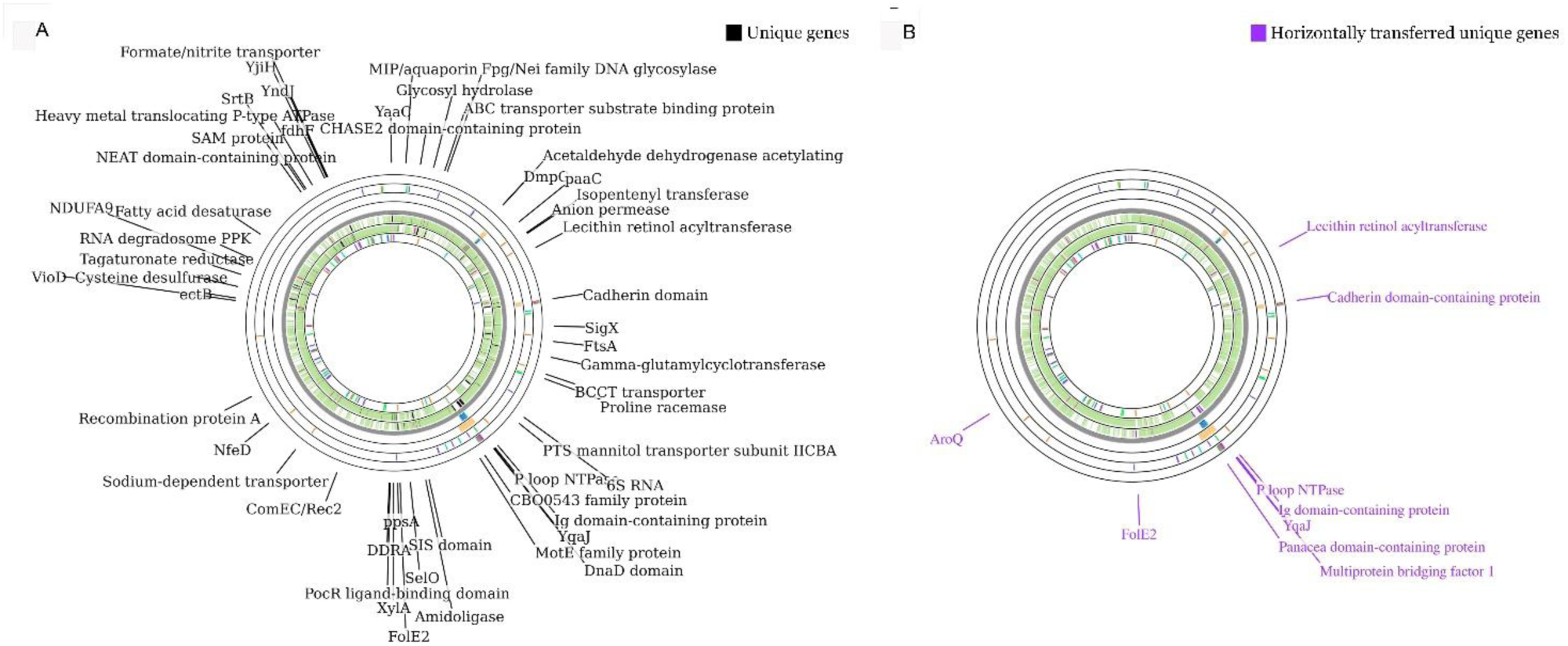
*Brevibacillus rhodotorulae* unique genes with known functions contain nine horizontally acquired genes. IslandViewer 4, Alien Hunter, and Alienness were used to identify Horizontally acquired unique genes in *Brevibacillus rhodotorulae* A. Circular genome representation of the functionally known unique genes of TJ4, B. Horizontally acquired genes among the unique genes of TJ4.

In prokaryotes, HGT occurs through conjugation, transformation, transduction, and the action of mobile genetic elements (MGEs). Prophages predicted by PHASTEST and HGT genes mapped by Proksee (Table 2, Fig 4B) only partially overlapped, suggesting that transduction is only one route of gene acquisition in *B*. *rhodotorulae*. MGEs predicted by mobileOG-db appear to contribute substantially to the acquisition of genes involved in functions such as transport, replication, and recombination (Table S3). Most transposases found in other *Brevibacillus* genomes were absent from the unique genes of *B*. *rhodotorulae*, except for a Mu transposase. Although HGT often contributes to genome expansion, *B*. *rhodotorulae* also showed signs of genome reduction, as reflected by its low CheckM value (82%). Unlike other *Brevibacillus* species analyzed, it lacks a CRISPR system. It also contains 113 predicted MGEs (Table S3), a relatively high number that supports frequent HGT and potential host adaptations.

**Table 2.**
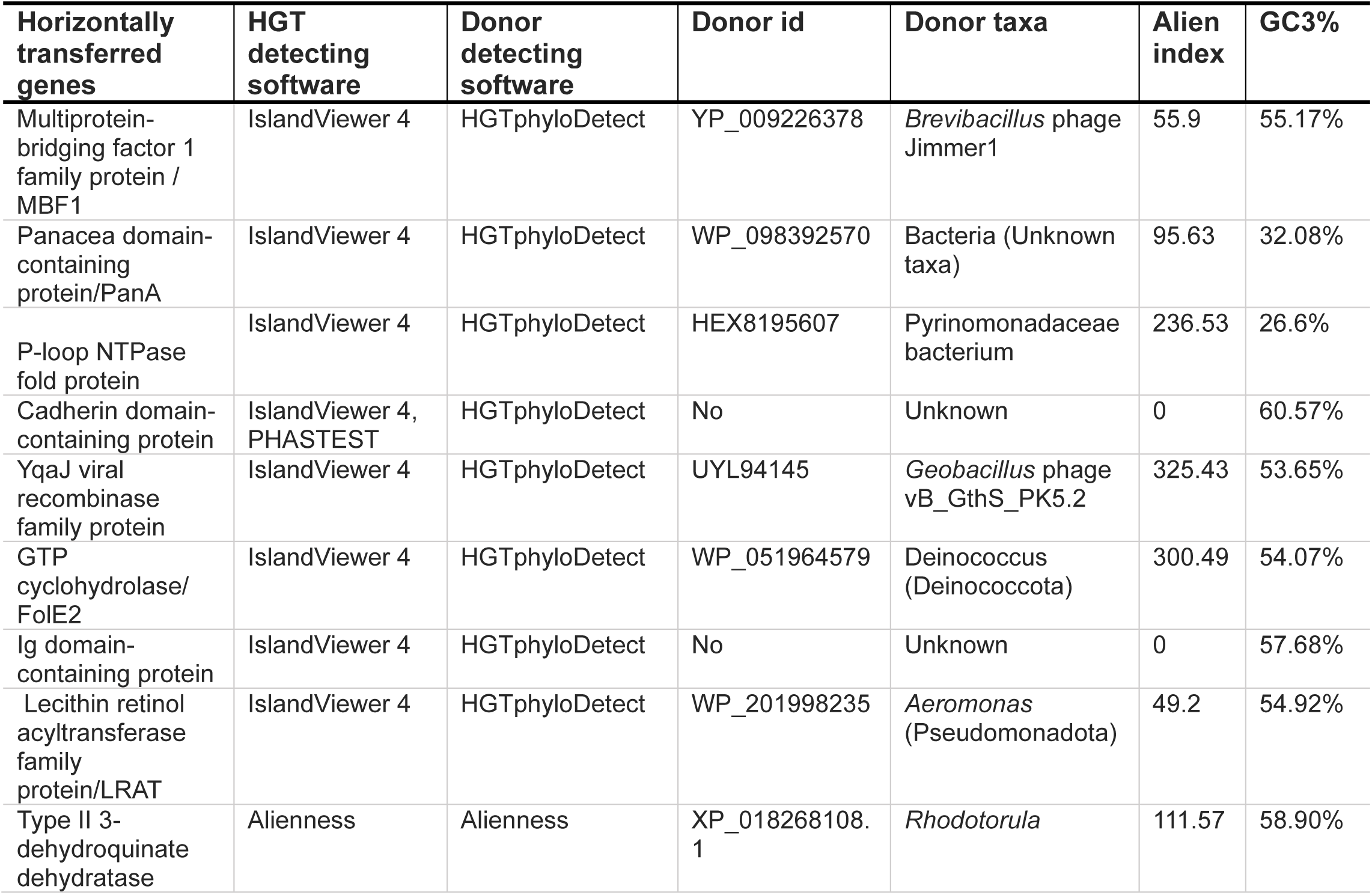
List of Horizontally transferred genes with potential donor organisms The HGT genes were analyzed using HGTphyloDetect to identify donor organisms. HGT from *Rhodotorula* was detected using Alienness.

#### JGTA-S1 to B. rhodotorulae gene transfer

Detecting HGT events in which eukaryotic genes are integrated into prokaryotes is particularly challenging because of their rarity. Alienness was used to study eukaryotic HGT events. A gene encoding a type II 3-dehydroquinate dehydratase (AroQ; Table 2), involved in the production of aromatic amino acids via the shikimate pathway, was acquired horizontally from *Rhodotorula* (Table 2). In *B*. *rhodotorulae*, AroQ is flanked by a sugar-phosphate isomerase/epimerase and a Bug family tripartite tricarboxylate transporter substrate-binding protein, which is closest to that of two distant *Brevibacillus* species (WP_023557546.1 and WP_280207186.1), respectively (Fig 5A). No homologous AroQ genes were detected in any *of the Brevibacillus* genomes closely related to *B. rhodotorulae*.

**Fig 5.**
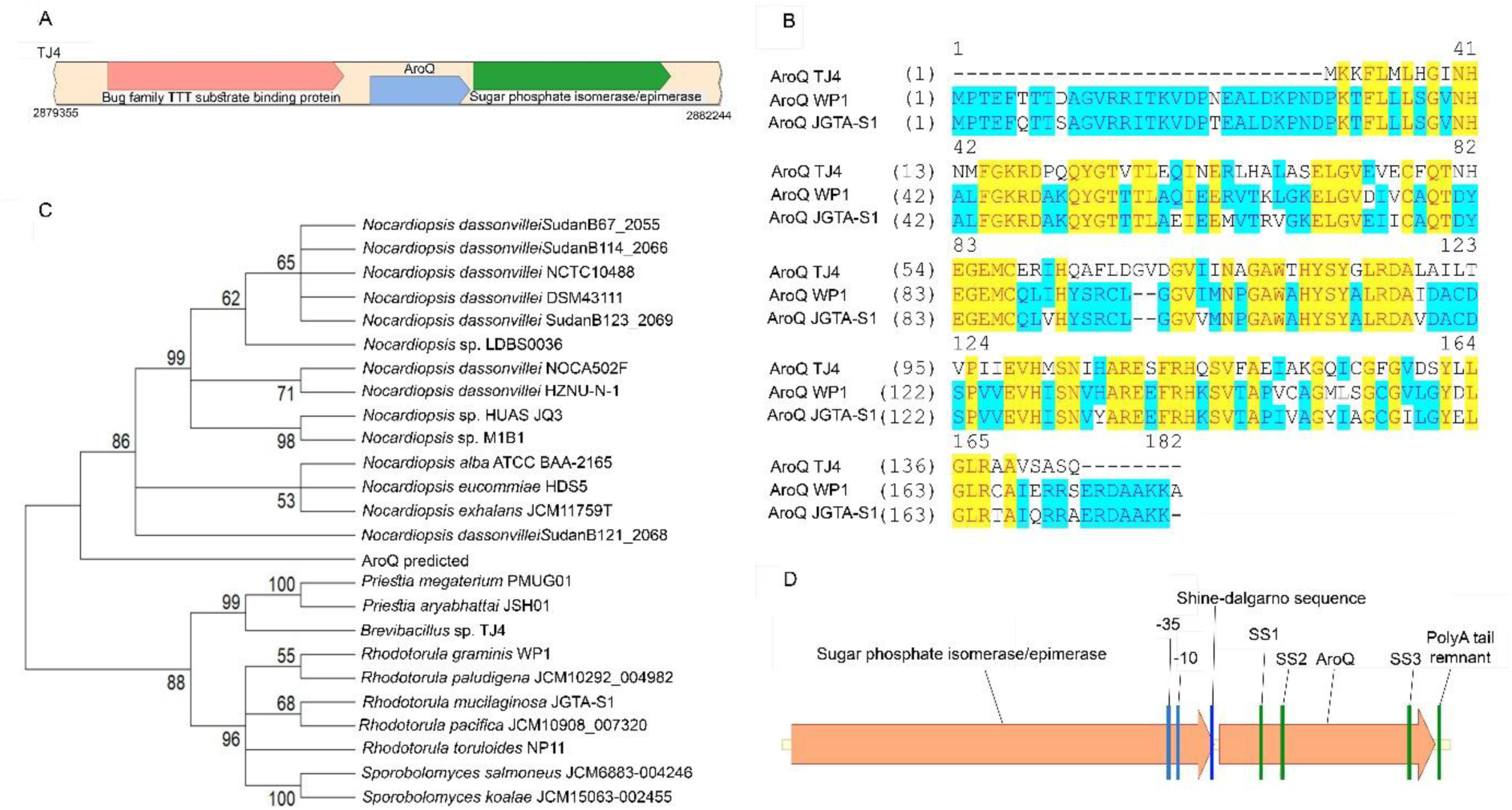
AroQ is horizontally acquired from *Rhodotorula* to TJ4. AroQ gene from JGTA-S1 and TJ4 were analyzed and compared to evaluate HGT of the fungal gene into *B. rhodotorulae*. A. SimpleSynteny was used to show the flanking genes of AroQ TJ4, B. Alignment of AroQ proteins from TJ4, *R. mucilaginosa* JGTA-S1, and *R. graminis* WP1, C. Gene Maximum Likelihood tree of the fungal and bacterial AroQ, D. Gene structure of AroQ TJ4 with eukaryotic features. SS = Splice site.

Reciprocal BLAST analyses were conducted using the AroQ TJ4 protein as a query against both fungi (taxid: 4751) and *the R. mucilaginosa* JGTA-S1 proteins used as local database. Additionally, we used JGTA-S1 AroQ as a query to search for bacterial sequences (taxid: 2). Reciprocal BLAST with AroQ TJ4 (WP_400164168.1) against fungi identified type II 3-dehydroquinate dehydratase from *R. glutinis* WP1 as the second-best hit. AroQ *B*. *rhodotorulae* returned JGTA-S1 004008-T1 (AroQ JGTA-S1) as the only hit in the JGTA-S1 proteome. Despite not being the first hit, querying with JGTA-S1 AroQ against bacterial sequences returned AroQ TJ4 with a significant e-value. The AroQ TJ4 protein showed substantial similarity to the JGTA-S1 and *R. glutinis* WP1 AroQ proteins (XP_018268108.1) (Fig 5B). We constructed a phylogenetic gene tree using the top fungal and bacterial gene hits obtained from reciprocal nucleotide BLAST analyses. Two CDS predicted from the JGTA-S1 genome by funannotate (JGTA-S1 004008-T1) and getorf (AroQ predicted) were included in the analyses (Fig 5C and Fig S1). AroQ TJ4 (*Brevibacillus* TJ4) nested within the fungal clade containing JGTA-S1-004008-T1 in the tree, suggesting that the genes share a common ancestor, potentially indicating HGT from the yeast to the bacteria. AroQ TJ4 has predicted - 10 and −35 consensus sequences 75 bp upstream of its ATG sequence, which might act as a functional promoter and a potential Shine-Dalgarno sequence (AGGAGT). In addition, AroQ TJ4 has three eukaryotic splice site motifs (GT-AG/GC-AG) and a poly A signal remnant (ATAAAAAA) (Fig 5D). All these findings support the HGT event of this AroQ gene from a close related yeast of JGTA-S1 to TJ4. AroQ is part of the shikimate pathway, which leads to the formation of chorismate, a key branch point in the synthesis of aromatic amino acids. The loss of modules in the shikimate pathway is not uncommon in host-associated bacteria that participate in mutualism and endosymbiosis (Zucko et al. 2010), suggesting that TJ4 acquired this essential component from its host, restoring at least part of the pathway that leads to chorismate and subsequent tryprophan production.

#### Horizontally acquired unique genes of B. rhodotorulae

Among the 84 unique genes identified in *B. rhodotorulae*, eight were predicted to have been acquired by horizontal gene transfer (Table 2): multiprotein-bridging factor 1 family protein (MBF1), panacea domain-containing protein, P-loop NTPase fold protein, cadherin domain-containing protein, YqaJ viral recombinase family protein, GTP cyclohydrolase FolE2, Ig domain-containing protein, and lecithin retinol acyltransferase family protein (LRAT) (Fig 4B). Their putative foreign origin was supported by GC3 analysis, which measures the GC content at the third codon position, a site that is typically under a weaker selective constraint. All eight genes showed substantially lower GC3 values (26.6%–60.57%) than the *B. rhodotorulae* core gene average (66.22%), consistent with their horizontal acquisition (Table 2).

To assess the potential donors of these eight genes HGTPhyloDetect was used. The predicted donors were *Brevibacillus* phages or bacteria, including at least two bacterial species, *Deinococcus* sp. and *Aeromonas* sp., which were identified as potential JGTA-S1 endosymbionts in whole-genome metagenomic data (Supplementary Table 1 of Nag et al., 2024) (Table 2). This finding suggests that *B. rhodotorulae* appeared to have acquired some of these genes from its community of co-resident endobacteria within JGTA-S1. However, the type II 3-dehydroquinate dehydratase attributed to *Rhodotorula* is not unique to TJ4 and is also present in *B*. *massiliensis* phR.

### The genome synteny in *B. rhodotorulae* seems to be disrupted

Loss of synteny around horizontally acquired genes is a hallmark of horizontal gene transfer (Sevillya et al., 2020). To examine this in *B. rhodotorulae*, six *Brevibacillus* genomes with no more than two contigs—*B*. *agri* DSM 6348 (DSM6348), *B*. *antibioticus* TGS2-1, *B*. *aydinogluensis* 4475, *B*. *borstelensis* CP19, *B*. *brevis* NCTC2611, and *B*. *parabrevis* B3 were selected for synteny analysis using SimpleSynteny. The genomes closest to *B. rhodotorulae*, *B*. *invocatus*, and *B*. *panachumi* were excluded because they were highly fragmented (>100 contigs) and contained no contigs with the target genes. Synteny was assessed at loci containing the eight horizontally acquired genes. The regions surrounding the cadherin domain-containing protein, FolE2, panacea domain-containing protein, and lecithin retinol acyltransferase showed complete rearrangement (Fig 6A–D). The P-loop NTPase fold protein, YqaJ, Ig domain-containing protein, and MBF1 were clustered within a 24,720 bp region in TJ4, but synteny in this region was strongly disrupted compared to other *Brevibacillus* genomes examined (Fig 6E).

**Fig 6.**
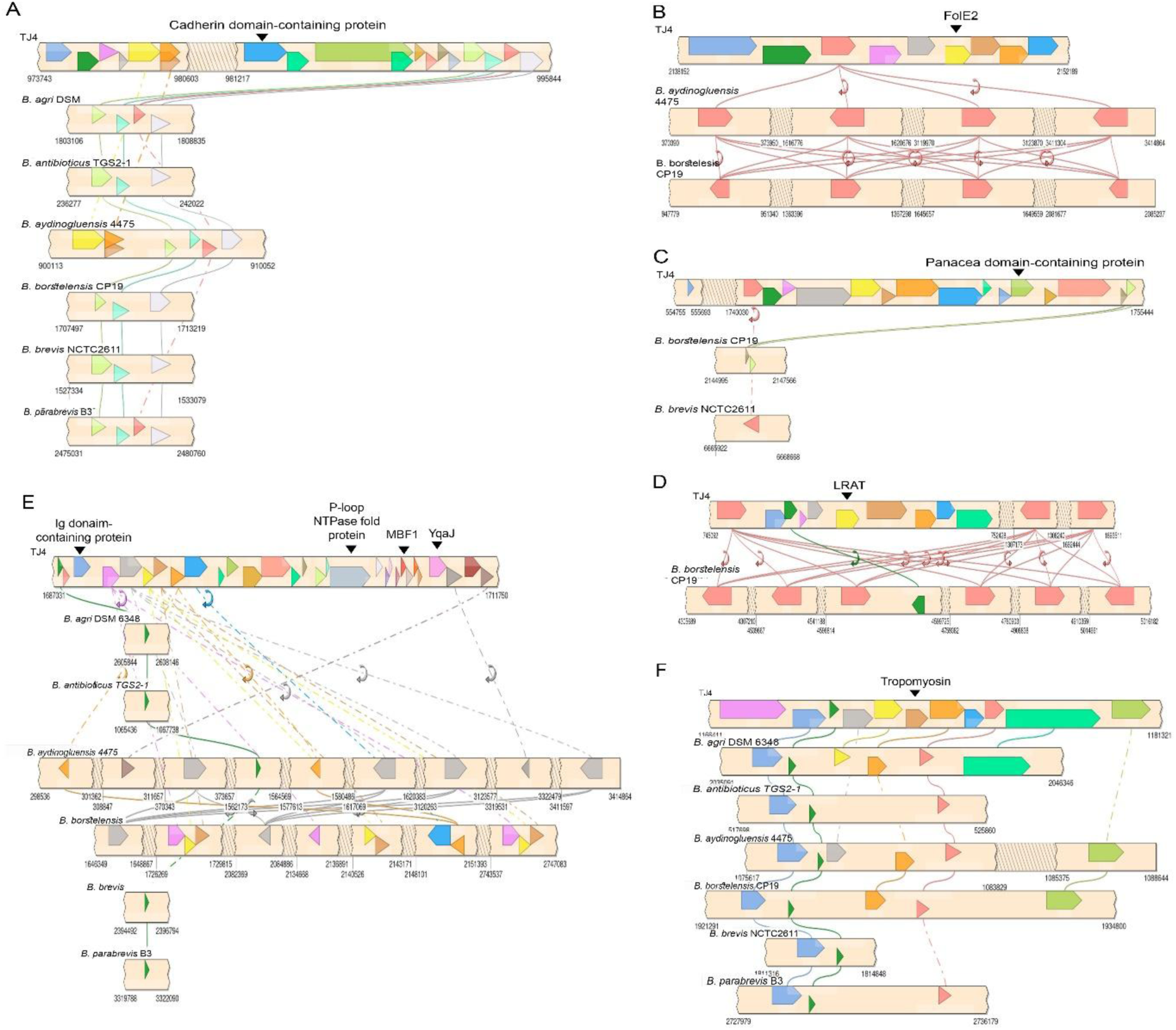
Rearrangement and or Loss of synteny in regions of TJ4 genome. SimpleSynteny was used to compare genomic regions of nine horizontally acquired genes from six *Brevibacillus* genomes A. Cadherin domain containing protein, B. GTP cyclohydrolase (FolE2), C. Panacea domain containing protein, lecithin retinol acyltransferase family protein (LRAT), E. Ig domain containing protein, P loop NTPase fold protein, MBF1, YQAJ, and F. Tropomyosin

A single-copy tropomyosin gene was identified in *B. rhodotorulae* between 5′-methylthioadenosine/adenosylhomocysteine nucleosidase and acetoin utilization protein (AcuC). Its flanking genes were conserved only in *B*. *agri* DSM 6348, and the disrupted synteny in other *Brevibacillus* species suggests the recent acquisition of this taxonomically restricted gene in *B. rhodotorulae* (Fig 6F). Tropomyosin, a component of the actin cytoskeleton, is widespread in eukaryotes and fungi and has only recently been reported in prokaryotes. In fungi and animals, it contributes to polarized growth, muscle contraction, and cytoskeletal dynamics. Tropomyosin is highly diverse in both eukaryotes and the bacterial genera *Streptomyces*, *Geobacillus*, *Pantoea*, *Paenibacillus*, *Bacillus* and *Microcystis*. In *B. rhodotorulae*, the predicted tropomyosin is a protein with a coiled-coil domain involved in chromosome segregation, potentially incorporated into the genome through illegitimate recombination facilitated by microhomologies.

## Discussion

Although *Brevibacillus rhodotorulae* can grow outside its host, its restricted capacity for independent survival mirrors traits of obligate endosymbionts, which typically undergo genome reduction and loss of key metabolic modules. Such reduction marks the transition from facultative to obligate symbiosis, as genes rendered redundant by host integration accumulate deleterious mutations and are eventually lost (Moran, 2002). At the obligate stage, endosymbionts possess highly reduced genomes, perfectly adapted to their hosts but incapable of free living. Small population sizes of endobacteria within JGTA-S1 further promotes genetic drift over natural selection, while long-term co-evolution stabilizes host–symbiont partnerships (Luong & Mathot, 2019).

The isolation of *B. rhodotorula*e has provided insights into the evolutionary trajectory of endosymbiosis. The majority of endosymbiotic microbes are categorized as either facultative or obligate. However, every obligate endosymbiotic relationship begins as a facultative relationship (Von Der Dunk et al. 2023), in which the interacting partners mutually benefit but can survive independently. *B. rhodotorula*e occupies a unique proto-obligate state, a transitional midpoint marked by an emerging metabolic dependency on the host. We use the term proto-obligate to describe the transitory phase of facultative endosymbionts to permanent host integration during the course of evolution. In the case of *B. rhodotorula*e, its dependence on its host is not yet absolute. Axenic cultivation of *B. rhodotorula*e is possible but remains challenging, for example, under stressful conditions, such as in minimal medium.

In contrast to the unicellular endosymbiosis observed in the marine alga *Braarudosphaera bigelowii*, which has a single cyanobacterial endosymbiont forming a nitrogen-fixing organelle known as the nitroplast (Coale et al., 2024), JGTA-S1 features many endosymbionts. This abundance suggests that endosymbiosis in JGTA-S1 is ongoing and relatively recent. However, the endosymbiotic relationship in *B. rhodotorulae* appears to be established enough for gene exchange, in which genes are gained or lost from its genome. CRISPR systems typically protect bacteria by preventing the entry of foreign DNA through adaptive immune responses (Wheatley and MacLean, 2021). The absence of a CRISPR system in *B. rhodotorulae* makes it more susceptible to foreign DNA invasion than other bacteria. There are two main reasons why its genome size is not as reduced as seen in other obligate endosymbionts. DNA loss common in early endosymbiosis seems to be offset by MGEs and HGTs. As *B. rhodotorulae* resides within the protective confines of yeast cells, many of its defense and protective mechanisms such as sporulation and CRISPR may have become dispensable.

Another observation was the high rate of HGT, including the potential transfer of an AroQ gene of the shikimate pathway in the *B. rhodotorulae* genome. In the absence of this gene, chorismate synthesis is impaired, preventing the bacterium from producing tryptophan, the only aromatic amino acid it can synthesize. Eukaryotic-to-prokaryotic HGT is rare because of the instability of eukaryotic genes in the prokaryotic genomic environment due to the absence of suitable promoters, lack of splicing mechanisms, and codon usage differences. This may result in premature termination, further complicating such integrations. However, recent evidence suggests that despite these obstacles, HGT from eukaryotes to prokaryotes occurs, especially when it results in a selective advantage for the recipient bacteria. Eukaryotic introns are modified inside bacteria, leading to the generation of functional proteins (Yuan et al. 2024). Two AroQ genes were predicted in JGTA-S1 from the positive and negative strands of the same region. The negative strand is predicted to code for an intron-containing gene JGTA-S1 004008-T1, but the positive strand is predicted to code for an intron-less gene (predicted AroQ). The phylogeny suggests that JGTA-S1 004008-T1 is closely related to *B. rhodotorulae* AroQ. AroQ in TJ4 has several features that are reminiscent of its eukaryotic origin. Yet it potentially possesses all the machinery required for transcription to occur in bacteria. Our evidence supports the hypothesis that the *B. rhodotorulae* AroQ gene originated from a yeast lineage closely related to that of JGTA-S1. A similar observation was made in the mycoplasma-related endosymbiont of arbuscular mycorrhiza *Dentiscutata heterogama*, where the horizontal acquisition of several host genes was detected in the bacteria (Torres-Cortés et al. 2015).

Several major metabolic pathways in *B. rhodotorulae* were lost under the influence of the yeast’s internal environment. Carbohydrate, amino acid, and vitamin metabolism are affected in other obligate endosymbionts, such as *Cytomitobacter indipagum, C. primus, Nesciobacter abundans, and Sneabacter namystus,* of the host Diplonemid (George et al. 2020). Endosymbionts often import essential nutrients from their hosts and cohabiting bacteria (Coale et al. 2024). *B. rhodotorulae* possesses 541 transporters and eight amino acid ABC transporter ATP-binding proteins, including those specific for lysine and arginine (for example, WP_400162589.1), as well as branched-chain amino acids such as isoleucine (eg. WP_421616994.1). This extensive repertoire of transporters potentially enables *B. rhodotorulae* to import amino acids that it cannot biosynthesize. Additionally, *B. rhodotorulae* contains 41 pseudogenes (Table S3), indicating evolutionary adaptation to the yeast host, which is also consistent with an endosymbiotic, slow-growing lifestyle. *Rhodotorula mucilaginosa* JGTA-S1 appears to function as a unique hub where endobacteria are constantly evolving, making it an ideal model for studying continuous endosymbiotic evolution.

## Methods

### Isolation of *Brevibacillus rhodotorulae*

*Rhodotorula mucilaginosa* JGTA-S1 was grown in a nitrogen-free medium (glucose, 5 g; yeast extract, 0.1 g; CaCl_2_,0.15 g; Na_2_MoO_4_, 0.005 g; MgSO_4_, 7H_2_O, 0.2 g; FeSO_4_,7H_2_O, 0.04 g; K_2_HPO_4_,0.5 g; CaCO_3_, 1.0 g; and distilled water to 1 L, pH 7.0) (Nag et al. 2024). The yeast cells were ruptured, and the cell extracts were plated on TSB (HiMedia, India) agar plates containing the antifungal agent amphotericin B (15 µg/mL) to inhibit the growth of JGTA-S1 yeast cells. The plates were incubated at 28°C for 3 d. Single yellow colonies were observed and streaked four times to obtain pure cultures. Glycerol stocks were then prepared.

### Genome sequencing, assembly, and annotation

Bacterial genomic DNA was isolated using the QIAamp DNA Mini Kit (Qiagen). The quality and quantity of the extracted DNA samples were checked using a Nanodrop spectrophotometer by determining the A_260_/A_280_ ratio, followed by agarose gel electrophoresis. Genomic DNA was sequenced using a combination of Illumina short-read (150 bp, paired-end) and Oxford Nanopore long-read sequencing (Oxford Nanopore PromethION P2 Solo). Sequencing was performed by Eurofins Genomics, and a hybrid de novo assembly was generated using SPAdes (v3.14.1) (Bankevich et al. 2012). Furthermore, the completeness of the assembly was checked using Benchmarking Universal Single-Copy Orthologs (BUSCO) and CheckM. Genes were predicted using Prokka (v1.12) (Seemann 2014), and functional annotations were performed using the Diamond tool in the BLASTX mode (E-value: 1e^-05^).

### Comparison of *B. rhodotorulae* with related *Brevibacillus* genomes

The Type (Strain) Genome Server (TYGS) (Meier-Kolthoff and Göker 2019) database and the Bacterial and Viral Bioinformatics Resource Center (BV-BRC, v3.54.6) were used for whole-genome-based taxonomic analyses. The TJ4 genome was compared with 17 other *Brevibacillus* genomes. These eighteen *Brevibacillus* species were *B*. *agri* DSM 6348 (GCA_004117055.1), *B*. *antibioticus* TGS2-1 (GCA_005217615.1), *B*. *aydinogluensis* 4475 (GCA_961514395.1), *B*. *brevis* NCTC2611 (GCA_900637055.1), *B*. *brevis* DSM 30 (GCA_003385915.1), *B*. *borstelensis* CP19 (GCA_036700025.1), *B*. *choshinensis* DSM 8552 (GCA_001420695.1), *B*. *composti* FJAT-54423 (GCA_016406105.1), *B*. *formosus* DSM 9885 (GCA_001012775.1), *B*. *fortis* NRRL NRS-1210 (GCA_003013395.1), *B*. *gelatini* DSM 100115 (GCA_003710935.1), *B*. *invocatus* JCM 12215 (GCA_003710915.1), *B*. *massiliensis* phR (GCA_000311785.1), *B*. *panacihumi* JCM 15085 (GCA_003710985.1), *B*. *parabrevis* B3 (GCA_022701015.1), *B*. *porteri* NRRL B-41110 (GCA_003013475.1), *B*. *reuszeri* DSM 9887 (GCA_001187725.1), and *Brevibacillus* sp. TJ4 (GCA_044627595.2).

### Pairwise comparisons of genome sequences

Digital DNA-DNA hybridization (dDDH) values and confidence intervals (CI) were calculated using the Genome-to-Genome Distance Calculator (GGDC) (Meier-Kolthoff et al. 2013, 2022). Dissimilarity in gene content (d0 and d6) and sequence identity in homologous regions (d4) were measured.

### Phylogenetic tree generation

A phylogenetic tree using 18 *Brevibacillus* genomes was generated using the MAFFT alignment tool with 100 unique genes and 40105 aligned amino acids using BV-BRC (Olson et al. 2023). RAxML Fast Bootstrapping was used as the branch support method. *Bacillus subtilis* subsp. subtilis str. 168 was used as an outgroup.

### Comparative genomics

Eighteen *Brevibacillus* genomes were used for comparative genomics using PanExplorer (Dereeper et al. 2022). *B*. *agri* DSM 6348, *B*. *antibioticus* TGS2-1, *B*. *aydinogluensis* 4475, *B*. *brevis* NCTC2611, *B*. *brevis* DSM 30, *B*. *borstelensis* CP19, *B*. *choshinensis* DSM 8552, *B*. *composti* FJAT-54423, *B*. *formosus* DSM 9885, *B*. *fortis* NRRL NRS-1210, *B*. *gelatini* DSM 100115, *B*. *invocatus* JCM 12215, *B*. *massiliensis* phR, *B*. *panacihumi* JCM 15085, *B*. *parabrevis* B3, *B*. *porteri* NRRL B-41110, *B*. *reuszeri* DSM 9887, and *Brevibacillus rhodotorulae* (GCA_044627595.2) were analyzed using PanACoTA to determine the core, dispensable, and unique genes. An UpSet plot was generated to identify overlaps between the functional clusters of the tested genomes. Unique TJ4 genes were represented using Proksee (Grant et al. 2023).

### Gene addition and deletion in *Brevibacillus rhodotorulae*

The Excel Power Query tool was used on the results from PanExplorer to identify the unique and lost genes in TJ4. KofamKOALA was used to allocate KEGG Orthologs (K numbers representing KOs) to protein sequences (E-value: 0.01) (Aramaki et al. 2020). KO assignments were sent to the KEGG Mapper reconstruction tool to identify the presence or absence of genes in each pathway.

### Detecting HGT events and donor taxa of horizontally transferred genes

Alien Hunter (Vernikos and Parkhill 2006), IslandViewer 4 (Bertelli et al. 2017), Alienness (Rancurel et al. 2017), and HGTphyloDetect (Yuan et al. 2023) software were used to detect HGT events in *B. rhodotorulae*. PHASTEST was used to detect prophages (Wishart et al. 2023). Mobile genetic elements were identified using mobileOG-db (Brown et al. 2022). Unique HGTs were represented using Proksee (Grant et al. 2023). Eight TJ4 protein sequences were analyzed using HGTphyloDetect to identify the donor of potential horizontal gene transfer (HGT) events. The protein sequences were subjected to a sequence similarity search using BLASTP via the blast.py script against the NCBI non-redundant (nr) protein database to identify homologous sequences from other taxa. The generated BLAST outputs were used as inputs for the HGTphyloDetect workflow, which was run using HGT_workflow_distant. py script. The Alien Index (AI=40) and outgroup percentage (out_pct=0.80) were used as default parameter thresholds to detect probable horizontally transferred genes. This pipeline generates tabulated results, such as Alien index values, E-values, donor identifiers, and corresponding donor taxonomic classifications. Alienness software was used to detect HGT genes in *B. rhodotorulae*. The protein sequences of TJ4 were queried against the NCBI nr database (nr_taxid.dmnd) using DIAMOND BLASTP with an e-value of 1e-^5^ threshold, retrieving up to 500 hits per query. The generated BLAST output file was uploaded to the Alienness web interface https://www.mdpi.com/2073-4425/8/10/248 against *Brevibacillus* sp. TJ4 taxonomic group (taxid: 3234853). Bacteria (taxid:2) and Archaea (taxid: 2157) were excluded from the analyses. The outputs were used to infer potential donors. AroQ TJ4 and AroQ JGTA-S1 were aligned with *R. graminis* WP1 (XP_018268108.1) using Vector NTI 10. Reciprocal BLASTP was performed using AroQ TJ4 and JGTA-S1 AroQ proteins against fungi (taxid: 4751) and bacteria (taxid: 2) sequences, respectively, to check for the presence of AroQ *Rhodotorula* sp. and AroQ TJ4 proteins among the top hits. Top fungal and bacterial gene sequences obtained by nucleotide BLAST of TJ4 and JGTA-S1 AroQ CDS were aligned using MEGA 12 with the MUSCLE algorithm. A phylogenetic tree was created using the Maximum Likelihood method with 1000 bootstrap replicates using the Kimura 2-parameter model. The AroQ TJ4 gene upstream region was searched for −10 and −35 using the BPROM tool (Cassiano and Silva-Rocha 2020). The gene was searched for eukaryotic splice site motifs (GT-AG/GC-AG), Shine-Dalgarno sequence (AGGAGT), and fragments of the polyA signal.

### Calculation of GC3% of horizontally acquired unique genes

Horizontally transferred genes were subjected to GC3% calculations using the Biopython SeqIO module.

### Loss of synteny analysis

SimpleSynteny (Veltri et al. 2016) was used to perform a loss-of-synteny analysis. Six closely related *Brevibacillus* species with no more than two contigs (*B. aydinogluensis* 4475, *B. brevis* NCTC2611, *B. borstelensis* CP19, *B. parabrevis* B3, *B. antibioticus* TGS2-1, and *B. agri* DSM 6348) were used as genome contig file. Upstream and downstream genes of nine unique horizontally transferred genes (based on AlienHunter, PHASTEST, and Alienness data) of TJ4 were uploaded as the gene query files. Loss of synteny was inferred through the absence of connections and changes in gene order and location.

### Oxidative stress assay

*Brevibacillus* sp. TJ4 and *Bacillus velezensis* TSB6.1 (accession no. NZ_JAOXLR000000000.1) were grown in TSB by incubating at 28°C with shaking at 150 rpm for 2 d. The bacterial cells were normalized in TSB at 1 OD_600_ and spotted (3 µL) on TSB plate containing 1 µM paraquat dichloride. Undiluted (OD_600_ 0.5), 1: 100 (OD_600_ 0.005), 1: 1000 (OD_600_ 0.0005).

### Auxotrophy assay

*B. rhodotorulae*, *B*. *laterosporus* (ATCC 64)*, B*. *parabrevis*, and *E*. *coli* were grown in minimal M9 (5X stock was prepared using 60 g Na_2_HPO_4_, 30 g KH_2_PO_4_, 5 g NaCl, 10 g NH_4_Cl, and distilled water to 1 L, pH 7). 1 L of M9 (1X) medium was supplemented with 2 mL of 1 M MgSO_4_, 20 mL of 20% glucose, and 0.1 mL of CaCl_2_. All bacterial cultures were initially grown in nutrient-rich TSB medium at 28°C with shaking at 150 rpm for 2 d. The cultures were centrifuged and washed three times with autoclaved 18.2 MΩ water to remove the remaining medium from the cell pellets. The cells were then suspended in M9 medium. Subsequently, all cell suspensions were normalized to an OD_600_ of 1. Hundred microlitres of the bacterial suspension was added as an inoculum in 5 mL of M9. The cultures were then grown at 28°C with shaking at 150 rpm for 14 d. Uninoculated M9 was used as a blank. The experiment was repeated twice.

### Dependency of *B*. *rhodotorulae* on JGTA-S1 extract

*B. rhodotorulae* was first grown in TSB and washed with 18.2 MΩ water and M9 (1X) medium. The bacterial cells were suspended in M9, and the OD_600_ was normalized to 1. Next, 100 μL of the suspension was subcultured in 5 mL of M9. JGTA-S1 was cultivated in Yeast Extract Peptone Dextrose (YPD) broth for 2 d at 28°C and 150 rpm. After ensuring the purity of the JGTA-S1 culture using microscopy, the JGTA-S1 cells were thoroughly washed and suspended in 18.2 MΩ water. The JGTA-S1 cells were mechanically broken using glass beads to prepare the extracts. Cell rupture was confirmed by microscopy. The cell extract was centrifuged, and the supernatant was passed through a 0.22-micron filter to obtain the JGTA-S1 extract without any cellular remnants. The total protein concentration of the extract was measured using Bradford’s reagent and was set to 100%. The TJ4 cells in M9 medium were supplemented with varying concentrations of JGTA-S1 extract (0, 2, 10, 20, and 30%). Uninoculated M9 was used as the negative control. All tubes were supplemented with amphotericin B (60 µg/mL). The cultures were then grown at 28°C with shaking at 150 rpm for 3 d. The experiment was performed in triplicate.

## Data access

All data generated or analyzed during this study are included in this published article [and its supplementary information files].

## Competing interest statement

The authors declare no conflict of interest

## Acknowledgements

We acknowledge the Ignite Life Science Foundation (Grant No.: AgSci/22-23/02) and the Science and Engineering Research Board (SERB), Department of Science and Technology, Government. of India (Grant number: CRG/2019/000378) for funding this work. AR thanks CSIR, India for his fellowship. We thank Professor Snezhana Oliferenko for her constructive feedback and insightful comments on this manuscript. We acknowledge Prof Arup Mitra for kindly providing us the *B. parabrevis* strain.

## Author contributions

TR Isolated the bacterium, performed the investigation, data curation, and formal analysis, and wrote the original draft of the manuscript. JP performed the investigation and data curation for Alienness and HGTphyloDetect. AR did the oxidative stress assay. AS acquired funding, conceptualized and supervised the study, and wrote, reviewed, and edited the manuscript.

## Notes

### Competing Interest Statement

The authors have declared no competing interest.

